# What is hidden in the darkness? Deep-learning assisted large-scale protein family curation uncovers novel protein families and folds

**DOI:** 10.1101/2023.03.14.532539

**Authors:** Janani Durairaj, Andrew M. Waterhouse, Toomas Mets, Tetiana Brodiazhenko, Minhal Abdullah, Gabriel Studer, Mehmet Akdel, Antonina Andreeva, Alex Bateman, Tanel Tenson, Vasili Hauryliuk, Torsten Schwede, Joana Pereira

## Abstract

Driven by the development and upscaling of fast genome sequencing and assembly pipelines, the number of protein-coding sequences deposited in public protein sequence databases is increasing exponentially. Recently, the dramatic success of deep learning-based approaches applied to protein structure prediction has done the same for protein structures. We are now entering a new era in protein sequence and structure annotation, with hundreds of millions of predicted protein structures made available through the AlphaFold database. These models cover most of the catalogued natural proteins, including those difficult to annotate for function or putative biological role based on standard, homology-based approaches. In this work, we quantified how much of such “dark matter” of the natural protein universe was structurally illuminated by AlphaFold2 and modelled this diversity as an interactive sequence similarity network that can be navigated at https://uniprot3d.org/atlas/AFDB90v4. In the process, we discovered multiple novel protein families by searching for novelties from sequence, structure, and semantic perspectives. We added a number of them to Pfam, and experimentally demonstrate that one of these belongs to a novel superfamily of translation-targeting toxin-antitoxin systems, TumE-TumA. This work highlights the role of large-scale, evolution-driven protein comparison efforts in combination with structural similarities, genomic context conservation, and deep-learning based function prediction tools for the identification of novel protein families, aiding not only annotation and classification efforts but also the curation and prioritisation of target proteins for experimental characterisation.

## Introduction

When we consider all possible sequences that can be built out of a simple alphabet of 20 amino acids, there is an infinite number of putative proteins. Since the sequencing of the first protein, large-scale genomic efforts brought about by the development of faster and cheaper genome sequencing efforts have shed light into some of the sequences that nature has sampled so far, with more than 200 million protein coding sequences deposited in UniProt and more than 2 billion in MGnify (Richardson et al., 2023; UniProt Consortium, 2023).

The rate by which this number is increasing is much faster than that by which each individual protein can be experimentally characterised. To close the gap, functional information is gathered for a subset of individual protein representatives and then the findings are extrapolated to close homologs that are identified by computational approaches. Manual curation of such functional annotation is carried out by those assembling the genomes and by biocurators (Boutet et al., 2016) most of which is incorporated into automated annotation pipelines such as InterPro (Paysan-Lafosse et al., 2023). However, despite the great success of such approaches, only 83% of UniProt sequences are covered by InterPro, and many of these correspond to Domain of Unknown Function (DUF) domains, indicating that there is a large number of protein sequences that remain functionally unannotated and unclassified. In 2009, it was predicted that circa 20% of the proteins deposited in sequence databases represented such “dark matter” of the protein universe (Levitt, 2009). Some of these proteins may just correspond to highly divergent forms of known protein families and thus lie beyond the detection horizon of the automated, homology-based methods employed; others could belong to so far undescribed protein families with yet-to-be determined molecular or biological functions.

The 3D structure of a protein is an important source of information about its molecular mechanism or biological role. Obtaining such data experimentally is, however, an expensive and time-consuming process. As of March 2023, only about 200’000 experimental 3D structures of proteins and protein complexes are known and made available through the Protein Data Bank (PDB) (Bittrich et al., 2023), which contrasts with the more than 320 million unique protein sequences in UniRef100 (Suzek et al., 2015). Computational protein structure prediction is a way of bridging this gap, but with the current wealth of experimentally determined protein structures, homology-based approaches can expand structural coverage only up to a certain point (Bienert et al., 2017). On one hand, there are not enough experimentally determined templates that cover the full wealth of sequenced proteins and, from the other, even when a remote homolog of known structure is available, the lower the sequence similarity the higher the uncertainty in the predicted model.

Deep-learning-based approaches have recently circumvented this limitation by achieving extremely high levels of accuracy for proteins without any known homologs of known structure. The main player here was AlphaFold2, a deep neural network that performs end-to-end protein structure prediction based on evolutionary, physical and geometric information (Jumper et al., 2021). Its success drove its developers at DeepMind, in collaboration with the EMBL-EBI, to produce predicted structural models for all natural proteins catalogued in the UniProt Knowledgebase (UniProtKB) and set up the AlphaFold database (AFDB) (Varadi et al., 2022). AFDB version 4 (AFDBv4) includes structural models for about 215 million protein sequences and excludes those from viruses and those with more than 1’300 residues (2’700 for selected proteomes). This means that it provides models not only for well-studied proteins whose 3D structure is unknown, but also for all of those that make the “dark matter” of protein sequence databases.

In this work, we combine sequence, structure and genomic context similarities with deep learning-based function prediction tools to shed light into these proteins and the reasons why some of them remain unannotated based solely on sequence similarity and homology-based approaches. For that, we first revised the proportion of such sequences in UniProtKB and constructed, for the first time, a sequence similarity network of all catalogued proteins with high quality predicted structural models. We then combined our results with protein structure information as predicted by AlphaFold2 and evaluated how much structural novelty is encompassed in these proteins, using an evaluation of substructure compositions based on natural language processing. We also looked into the diversity of protein names, both those derived from homology transfer and those predicted by recent deep learning methods.

Our analysis demonstrates that functional annotation of proteins, even from a purely computational perspective, requires a combination of data sources and approaches. We show this with four examples of cases where (1) while neither sequence nor structure similarity could pinpoint a possible protein function, analysis of genomic context and looking for remote homologs of genomic neighbourhood provided an experimentally testable hypothesis, resulting in the definition of a novel toxin-antitoxin system superfamily TumE-TumA; (2) structure alignment and remote homology searches find novel subfamilies; (3) deep-learning based protein name prediction gives a variety of names for a set of related proteins that form previously undefined protein families; and (4) examining structural outliers finds novel fold families and incorrect or incomplete proteins.

## Results and Discussion

### 1. Functional darkness in the UniProt and AlphaFold databases

As of August 2022, there were more than 520 million unique protein sequences in UniProt (i.e., UniRef100 clusters). This includes more than 500’000 manually curated proteins in Swiss-Prot, more than 220 million sequences in TrEMBL and more than 300 million sequences in UniParc from other sources not yet included in UniProtKB. The total number of sequences drops drastically to circa 50 million unique sequences if one filters the UniRef100 dataset to a maximum sequence identity of 50% (UniRef50). We define the “functional brightness” of a given protein as the full-length coverage with annotations of its close homologs, with 0% meaning “dark” and 100% meaning “bright” and a UniRef50 cluster being as “bright” as the “brightest” sequence it encompasses (Fig. 1A). For that, we only considered those annotations that correspond to domains and families whose title does not include “Putative”, “Hypothetical”, “Uncharacterised” and “DUF”, and included those corresponding to structural features as coiled coil segments and predicted intrinsically disordered stretches. With this, we excluded any functional darkness resulting from predicted intrinsically disordered or coiled-coiled proteins, and focused solely on that from proteins with a potential for a globular (or other) fold type.

**Figure 1.**
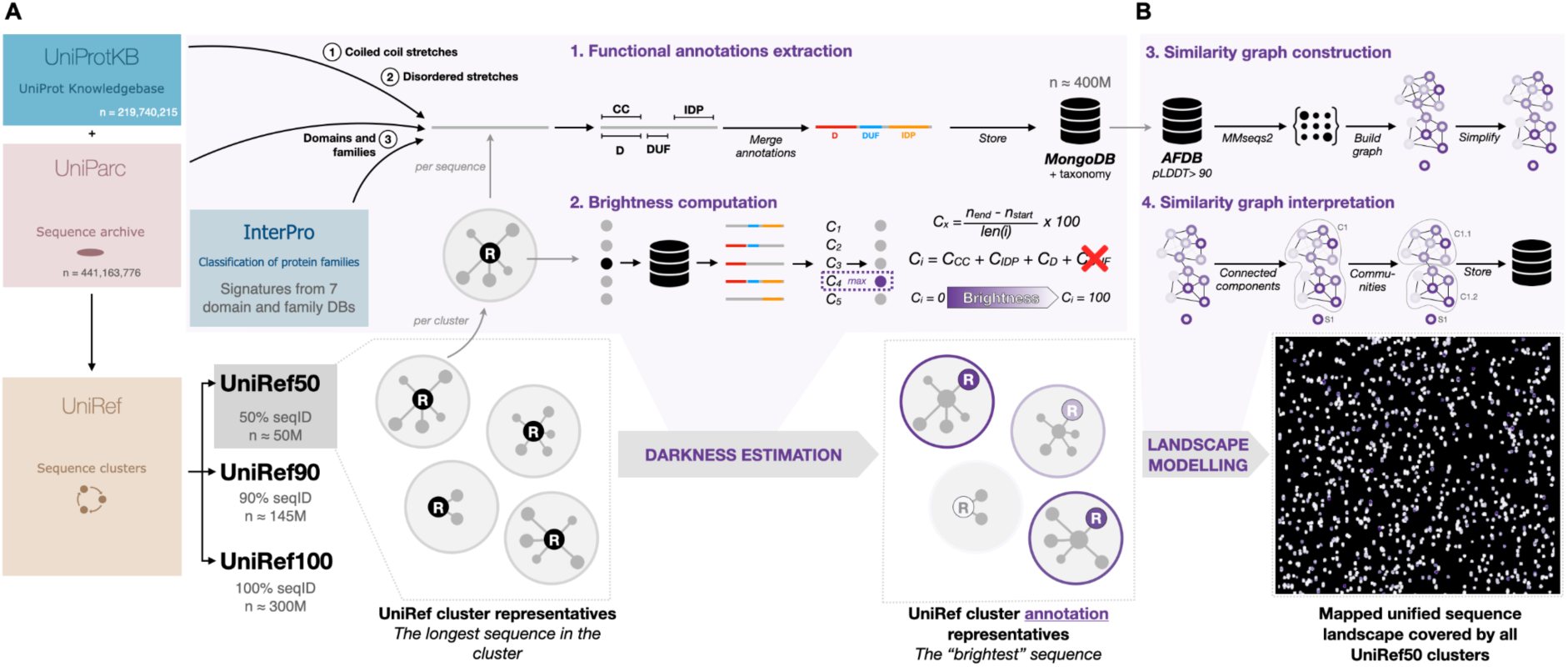
General workflow for the collection, classification and mapping of functionally dark proteins in UniProt and AlphaFold database. (A) Starting from the clusters in UniRef50, we collected all the functional annotations for all included UniProtKB and UniParc entries, excluding all of those with “Putative”, “Hypothetical”, “Uncharacterised” and “DUF” in their names, and selected as the functional representative of each cluster the protein with the highest full-length annotation coverage (i.e., brightness). (B) From the collected UniRef50 clusters, we selected only those with a structural representative with pLDDT >90, and constructed a large-scale sequence similarity network by all-against-all MMseqs2 searches, representing the sequence landscape of more than 6 million UniRef50 clusters.

By looking at the functional brightness distribution across all UniRef50 clusters, we observed that 34% of them do not reach a value higher than 5% (Fig. 2A). This means, 17’878’697 UniRef50 clusters have none of their protein sequences annotated for domains and families of known function, or for disorder and coiled coil propensities. All together, these clusters encompass 37’761’109 sequences, corresponding to 10% of all UniRef100. Strikingly, only 0.09% of them correspond to clusters of proteins assigned to a predefined DUF family (Fig. 2E).

**Figure 2.**
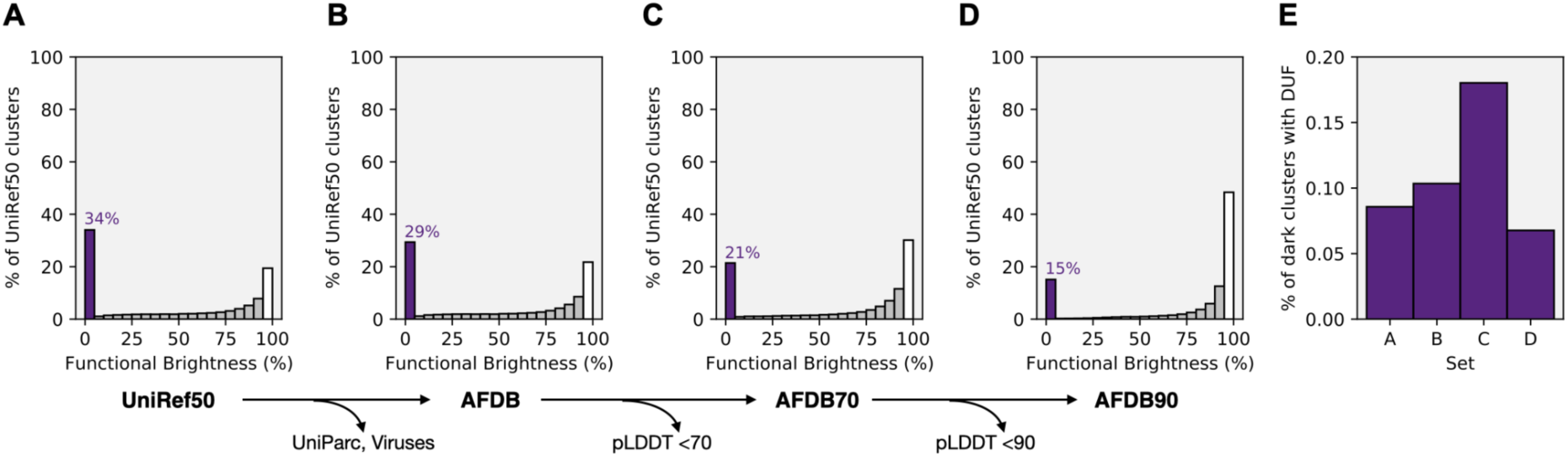
Distribution of functional darkness in UniProt and AFDB. Functional brightness distribution in (A) UniRef50, (B) UniRef50 clusters with models in AFDB (which excludes long proteins, and those UniRef50 clusters composed solely of UniParc entries and viral proteins), (C) UniRef50 clusters whose best structural representative has an average pLDDT > 70, and (D) UniRef50 clusters whose best structural representative has an average pLDDT > 90. For each set, the percentage of fully dark UniRef50 clusters, and corresponding brightness bin, are highlighted in purple. The bar associated with functionally bright UniRef50 clusters (functional brightness >95%) is marked in white. (E) Percentage of fully dark UniRef50 clusters with proteins annotated as a domain of unknown function (DUF) in each set A-D.

While the functional brightness of a UniRef50 cluster is not directly proportional to the number of sequences it groups (Pearson correlation coefficient of 0.0), fully bright clusters (functional brightness ≥ 95%) tend to be larger than those whose members are poorly annotated, with an average of 19 ± 123 non-redundant sequences. Indeed, fully dark UniRef50 clusters are composed, on average, of only 2 non-redundant sequences, but cases with more than 2’000 sequences were also found (e.g. UniRef50_A0A091P6G5, which corresponds to multiple transposons).

Thanks to AFDBv4, we now have access to high quality models for most of the proteins in UniProtKB (excluding UniParc and viruses). We observed that only 41’983’943 UniRef50 clusters (78% of all clusters) have members with a predicted structure in AFDBv4, and that 29% of these are functionally dark (Fig. 2B). The percentage of dark UniRef50 clusters decreases when considering only those UniRef50 clusters with high and extremely high predicted accuracy models (Fig. 2C,D), with 21% dark clusters with a structural representative with an average pLDDT > 70, and 15% with an average pLDDT > 90. This highlights that (1) there is a considerable amount of dark UniRef50 clusters without models in AFDB, (2) but there is also a good quantity of functionally unknown proteins for which there is structural information that we can now use to learn about them.

We observe that the proportion of DUFs in UniRef50 and AFDBv4 remains in the range of 0.1-0.2% independently of the pLDDT cut-off (Fig. 2E). This indicates that most dark UniRef50 clusters are not associated with a previously defined DUF, and AFDBv4 harbours high quality structure information for many more, yet to be defined, families.

### 2. Large-scale sequence similarity network of AFDB90

For the remainder of the work, we focused only on those dark UniRef50 clusters with high average pLDDT representatives, as these represent protein clusters that can be more confidently studied at a structural level. While UniRef50 clustering provides us with groups of sequences that are overall identical at the sequence level, such clustering does not reach the family and superfamily levels and does not account for local similarities. Thus, to search for those UniRef50 clusters that belong to the same families and superfamilies and identify those that are locally identical, we constructed a large-scale sequence similarity network of all Uniref50 clusters whose structural representative has an average pLDDT > 90 (the AFDB90 dataset). This corresponds to 6’136’321 UniRef50 clusters and, thus, to only 11% of the initial set of UniRef50 clusters.

To construct the network in a time efficient manner, we employed MMseqs2 (Steinegger and Söding, 2017) for all-against-all sequence searches (Fig. 1B). In these searches, two sequences were deemed as adjacent if there was an alignment that covered at least 50% of one of the proteins at an E-value better than 1×10^−4^. After simplification of the resulting graph by considering the 4 nearest neighbours of each node, we obtained a graph with 10’339’158 edges involving 4’270’404 nodes, which include 43% of the dark UniRef50 clusters in the full dataset (Fig. 3). The resulting graph (Fig. 3A, https://uniprot3d.org/atlas/AFDB90v4) is composed of 242’876 connected components with at least 2 nodes, with the largest encompassing about 50% of all UniRef50 clusters. Out of these components, 46’318 (19%) have an average brightness content below 5% and are, thus, “fully dark galaxies” in the protein sequence space represented by the network (Fig. 3D). However, not all dark UniRef50 clusters are enclosed in these dark connected components. Indeed, these components count only for 241’203 (60%) of the dark UniRef50 clusters in the graph. The remaining 40% are spread in components of variable brightness and thus represent remote global or local homologs of well- or partially well-annotated proteins.

**Figure 3.**
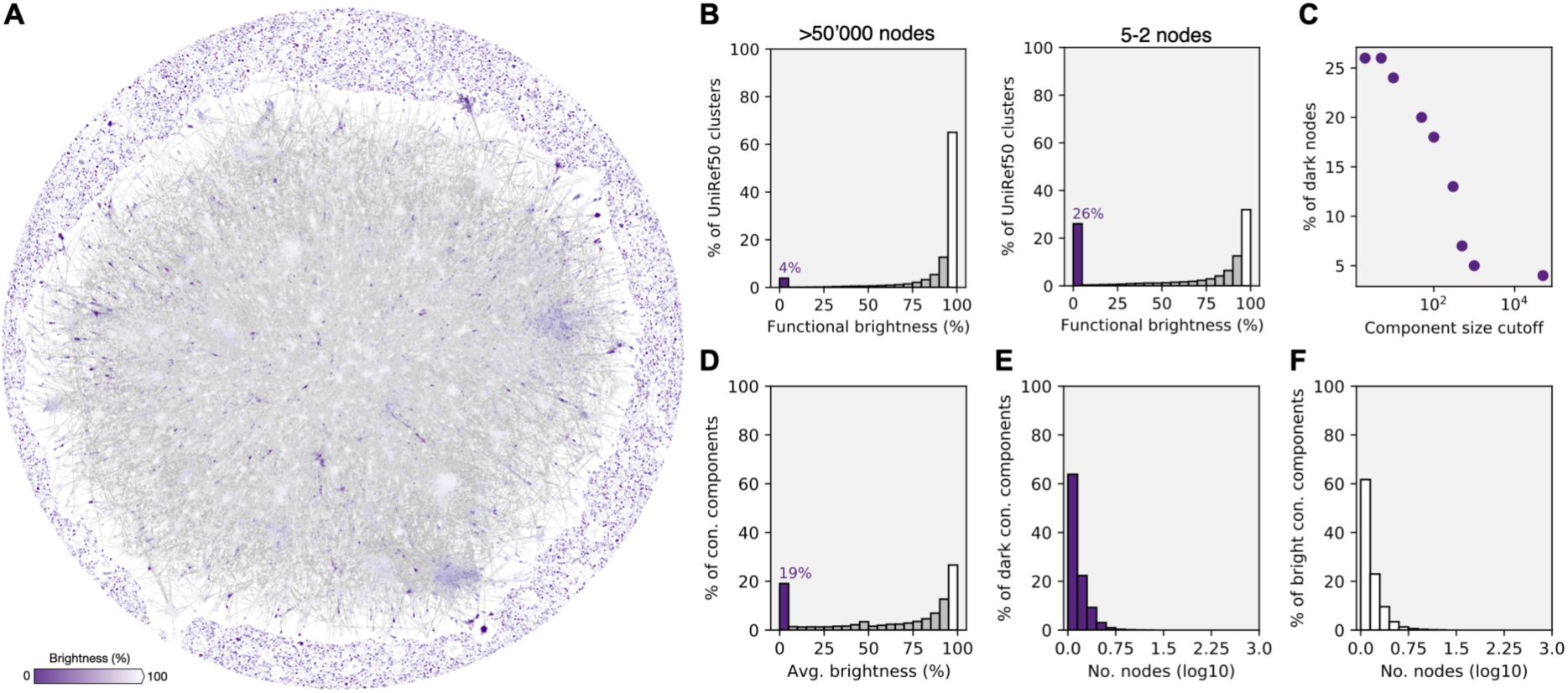
Large-scale sequence similarity network for over 6 million UniRef50 cluster representatives with high predicted accuracy models in AFDB (AFDB90). (A) Layout of the resulting network, as computed with Cosmograph (https://cosmograph.app/). For simplicity, the network was reduced to a set of 688’852 communities connected by a total of 1’488’764 edges. The 1’865’917 UniRef50 clusters that did not connect to any other in the MMseqs2 searches were excluded. Only the 473’612 communities that have at least one inbound or outbound edge (degree of 1) are displayed in the figure. Nodes are coloured by the average functional brightness of the UniRef50 clusters included in the corresponding community. An interactive version is available at https://uniprot3d.org/atlas/AFDB90v4, where all singletons are arranged outside the main circle. (B) Histograms of functional brightness content for connected components with more than 50’000 and with only 5 to 2 nodes (UniRef50 clusters), highlighting their different darkness content. (C) Scatter plot of the component size (i.e. number of UniRef50 clusters) cut-off and the percentage of functionally dark UniRef50 clusters. (D) Distribution of the average brightness per component. Size distribution for (E) fully dark connected components (average brightness < 5%) and (F) fully bright connected components (average brightness > 95%).

As expected, the percentage of UniRef50 clusters in dark connected components decreases with the component’s size (Fig. 3B,C), meaning that the lower the number of homologs the harder a protein is to annotate. Still, and while the distribution is skewed towards smaller sizes in both fully dark (i.e., average functional brightness <5%) and fully bright (i.e., average functional brightness >95%) connected components (Fig. 3E,F), the largest dark connected component in our network has 836 members, totalling 4’889 unique proteins. We selected two examples for further analysis: (1) component 27, which is the largest functionally dark connected component; and (2) component 159, a large component composed of various prokaryotic proteins without any structural homolog in the PDB.

### 3.1. A new family in a well-studied superfamily of transmembrane glycosyltransferases

The largest functionally dark connected component in our set is component 27, with 836 UniRef50 entries that include a total of 4’889 unique bacterial protein sequences (Fig. 4A). The structural representatives for each UniRef50 cluster have a median length of 665 ± 169 amino acids, and most are predicted to be transmembrane. Foldseek (van Kempen et al., 2022) finds multiple, medium confidence (TM-score ∼0.6) matches in the PDB for the structural representative with a length closest to the median (UniProtID A0A7X7MB17, Fig. 4B). These matches include multiple structures of eukaryotic Dolichyl-diphosphooligosaccharide-protein glycosyltransferase subunit STT3 and its bacterial homolog oligosaccharyltransferase PglB (Kelleher and Gilmore, 2006; Szymanski and Wren, 2005), both of which are absent from our network because their structure representatives have an average pLDDT < 90. Indeed, the proteins in this component that are not called “Uncharacterised protein” mostly have the title “YfhO family protein”, which corresponds to a family important for lipoteichoic acid or wall teichoic acid glycosylation (Rismondo et al., 2018). However, the representative of component 27 superposed poorly to the representative of the YfhO family (TM-score 0.58) and the average brightness of this component is 2±13%, with only 9 UniRef50 members (corresponding to 101 unique protein sequences) being full-length annotated as containing the “YfhO” InterPro domain.

**Figure 4.**
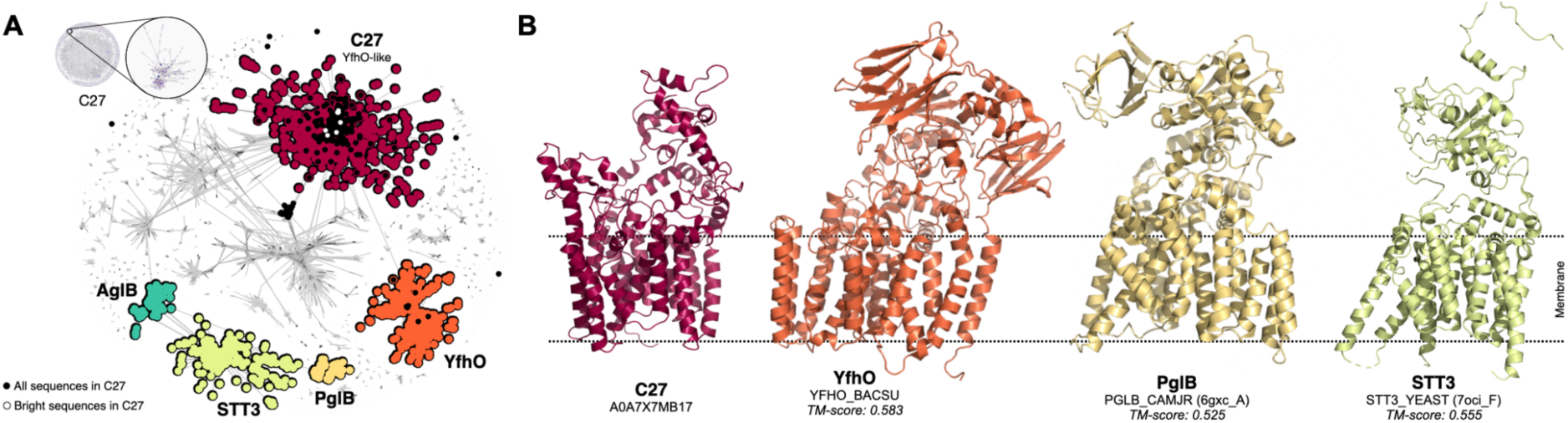
Connected component 27 is a new family in a well-studied superfamily of transmembrane glycosyltransferases. (A) High resolution sequence similarity network for 7’004 homologs of the sequences in component 27, computed with CLANS at an E-value threshold of 1×10^−20^. Points represent individual proteins and grey lines BLASTp matches at an E-value better than 1×10^−20^. Individual clusters are coloured and labelled accordingly to their representative members. Only YfhO-like and STT3/PglB sequences are highlighted, with grey dots depicting other homologous groups. AglB corresponds to the PglB/STT3-like sequences from archaea. Black dots depict those sequences that make component 27 in our network, and white dots mark those that are bright. (B) Predicted structural models as in AFDBv4 for the representative of component 27 (C27), and YfhO, and experimental structures of the PglB and STT3 cluster representatives. Models are coloured according to the colour of their corresponding cluster in (A). The membrane regions, as predicted with PPM 3.0 server (Lomize et al., 2022), are marked by dashed lines.

In order to understand the relationship between those proteins in component 27, STT3, PglB and YfhO, we combined the sequences of all four groups and built a sequence similarity network (Fig. 4A). This network highlights that most dark proteins in component 27 are not members of the same group as the reference YfhO and make a cluster by themselves. The YfhO-like protein family is linked to the STT3/PglB groups by multiple hypothetical proteins, mostly of prokaryotic origin, that are in some cases annotated as “Glycosyltransferase family 39 protein”. These results indicate that component 27 is a hitherto undescribed bacterial protein family belonging to a well-studied superfamily of transmembrane oligosaccharyl- and glycosyltransferases and may also be involved in the glycosylation of membrane lipids. This example illustrates the power of structural alignment to resolve undetected remote homology relationships in this new age of widespread predicted protein structure availability.

### 3.2. A new toxin-antitoxin superfamily

Component 159 (Fig. 5) is composed of 327 UniRef50 clusters, which correspond to 1222 unique protein sequences, whose members are mostly annotated as “Domain of Unknown Function 6516” (i.e. DUF6516). According to AFDB, these proteins may adopt a conserved α+β fold, where two α-helices pack against an antiparallel β-sheet with 7 strands (Figs. S1). Contrary to component 27, structural similarity searches over the PDB with a structure representative (UniProtID A0A6N7ITV5) using Foldseek found no matches at a TM-score better than 0.5.

**Figure 5.**
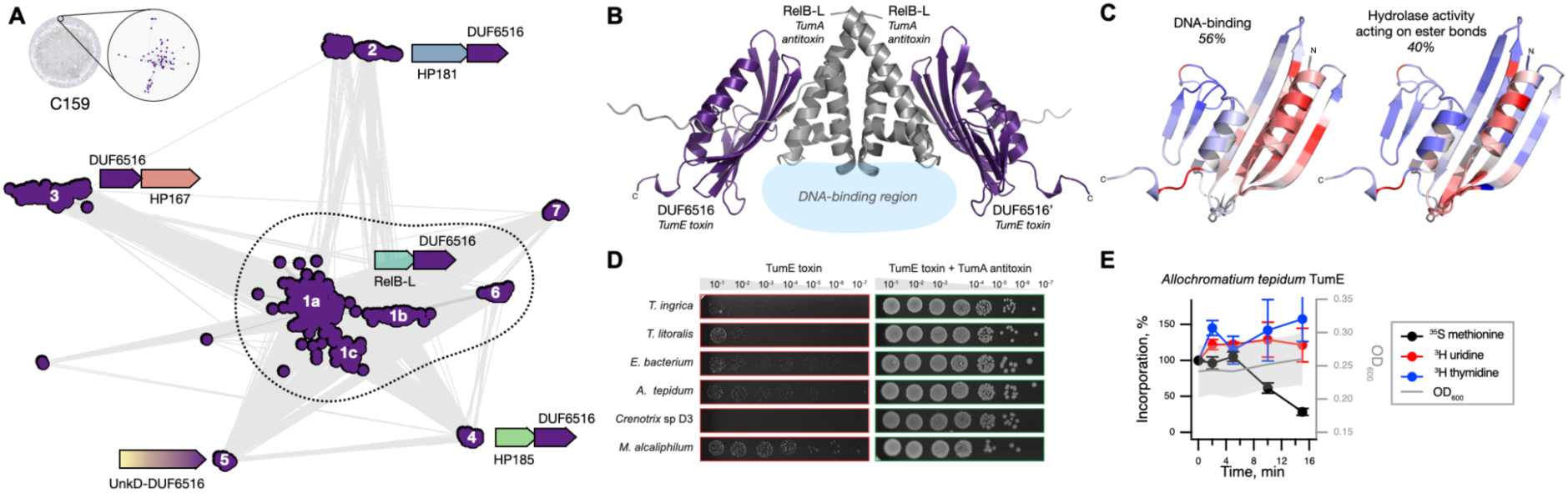
Connected component 159 is a novel toxin in the hitherto undescribed toxin-antitoxin superfamily TumE-TumA. (A) High resolution sequence similarity network for 2’453 homologs of the 327 sequences in component 159, computed with CLANS at an E-value threshold of 1×10-10. Points represent individual proteins and grey lines BLASTp matches at an E-value better than 1×10-4. Individual subclusters are labelled 1-7, with their subclusters labelled a-c. The consensus genomic context as identified with GCsnap (Pereira, 2021), with different flanking families coloured from blue to red and labelled accordingly. A gradient fill highlights the fusion between DUF6516-encoding genes and their putative antitoxin (which occurs in cluster 5, where the corresponding proteins are fused with a domain of unknown function and do not occur in a conserved genomic context). (B) 3D model of the complex between the putative toxin and antitoxin from *Allochromatium tepidum strain NZ*, modelled with AlphaFold-Multimer, highlighting the regions where DNA would interact with the antitoxin based on DeepFRI predictions and the structural features predicted for the antitoxin that resemble DNA binding regions. (C) Structural model of *A. tepidum* TumE/DUF6516 toxin (EntrezID WP_213381069.1) coloured according to the two most frequent molecular functions predicted for 100 homologs with DeepFRI. The corresponding residues responsible for the predictions are highlighted in red, and the percentage labelled reflects the frequency of the highlighted prediction. (D) Validation of tumE-tumA TA pairs using toxicity neutralization assay. Putative toxin expression plasmids (pBAD33 derivates) were cotransformed into *E. coli* BW25113 cells with cognate antitoxin expression plasmids or the empty pMG25 vector. Bacteria were grown for five hours in liquid LB media supplemented with appropriate antibiotics and 0.2% glucose for suppressing the production of putative toxins. The cultures were normalized to OD600 = 1.0, serially diluted and spotted on LB agar plates containing appropriate antibiotics and 0.2% arabinose for toxin induction and 500 µM IPTG for antitoxin induction. The plates were scored after an overnight incubation at 37 °C. (E) Metabolic labelling assays with wild-type *E. coli* BW25113 expressing *A. tepidum* TumE/DUF6516 toxin.

We constructed a high resolution similarity network for component 159 representatives, additionally enriched with close sequence homologs (Fig. 5A). The network contains 7 distinct classes of DUF6516-containing proteins, which DeepFRI (Gligorijević et al., 2021) predicted may bind DNA or other nucleic acids and carry a hypothetical catalytic site with a potential hydrolase activity over ester bonds (Fig. 5C, Supplementary file 1), To gain insight into the potential biological functions of DUF6516, we used the high-resolution network as input for GCsnap (Pereira, 2021). This tool compares the genomic contexts of the target homologous protein-coding genes and annotates the neighbourhood outputs with functional and structural information retrieved from UniProt and the SWISS-MODEL repository (SMR) (Bienert et al., 2017), respectively. Strikingly, DUF6516 is commonly found in a conserved two-gene (bicistronic) genomic arrangement, with DUF6516 predominantly located downstream of the conserved bicistronic “partner” (clusters 1, 2, 4 and 6).

While most of the “partner” genes associated with DUF6516 code for “hypothetical proteins” of unknown function, one of the partners in cluster 1 is a remote homolog of RelB, a well-characterised antitoxin (Gotfredsen and Gerdes, 1998). The bicistronic arrangement is typical for toxin-antitoxin (TA) systems (Jurėnas et al., 2022). When active, the TA toxin proteins abolish bacterial growth, and the control of this toxicity is executed by the antitoxin, which, in the case of so-called “type II TA systems”, is a protein that acts by forming an inactive complex with the toxin. Structure-based molecular function prediction for DUF6516 partners using DeepFRI (Gligorijević et al., 2021) suggests they may bind DNA (Supplementary file 1), an activity characteristic for diverse antitoxins (Jurėnas et al., 2022), and co-folding prediction with AlphaFold-Multimer generated high confidence models (93 average pLDDT, 0.902 iPTM) that support the interaction between the two proteins as a dimer of dimers (Fig. 5B), as observed for type II TA systems. Therefore, we hypothesised that DUF6516 is a novel toxic TA effector that is neutralised either *in trans* by diverse unrelated antitoxins (subclusters 1-4, 6 and 7) or *in cis* by a fused unknown antitoxin domain (UnkD, subcluster 5).

To validate the putative TAs experimentally and gain insights into the mechanism of DUF6516-mediated toxicity, we used our established toolbox for TA studies (Kurata et al., 2022). We targeted TA from six Gammaproteobacterial species (*Thioploca ingrica, Thiothrix litoralis, Crenothrix sp. D3, Methylotuvimicrobium alcaliphilum, Ectothiorhodospiraceae bacterium, Allochromatium tepidum* strain NZ) for testing in *E. coli* surrogate host, and all the putative toxins dramatically abrogated *E. coli* growth (Fig. 5D) while the putative antitoxins had no effect (Fig. S2). Neutralisation assays showed full suppression of toxicity when the toxins were co-expressed with cognate antitoxins (Fig. 5D), thus directly validating that these gene pairs are, indeed, *bona fide* TA systems. To probe the mechanism of DUF6516-mediated toxicity, we carried out metabolic labelling assays with ^35^S methionine (a proxy for translation), or ^3^H uridine (a proxy for transcription) or ^3^H thymidine (a proxy for replication). Expression of *Allochromatium tepidum strain NZ* DUF6516 toxin resulted in a decrease in efficiency of ^35^S methionine incorporation (Fig. 5E), indicative of the inhibition of protein synthesis. We hypothesize that the effect could be mediated by the yet-unproven RNase activity of the DUF6516 toxin. We conclude that DUF6516 is a *bona fide* translation-targeting toxic effector of a novel TA family, and propose renaming it TumE (for “dark” in Estonian), with the antitoxin components dubbed as TumA, with A for “antitoxin”.

This example illustrates the difficulty of automating functional annotation for proteins from completely novel superfamilies without any functionally characterised member. In this case, the combination of genomic context information, remote homology searches on genomic neighbours, and deep learning-based function prediction helped formulate a functional, testable hypothesis.

### 3. Semantic diversity of protein names in functionally dark connected components

Recently, ProtNLM (Gane et al., 2022), a large language model, was implemented as an approach to automatically name proteins in TrEMBL titled as “Uncharacterised protein”. Upon the first release of such predictions in UniProtKB, we investigated how diverse the names predicted for those proteins in dark connected components were, and compared it to those fully bright components. In both cases, we observed that the distributions of name and word diversities (collectively referred to as “semantic diversity”) across components were highly skewed towards extremely low values, but the fully dark set was significantly different in both cases from the fully bright set (Kolmogorov–Smirnov *p-*value = 1×10^− 16^, Fig. 6).

**Figure 6.**
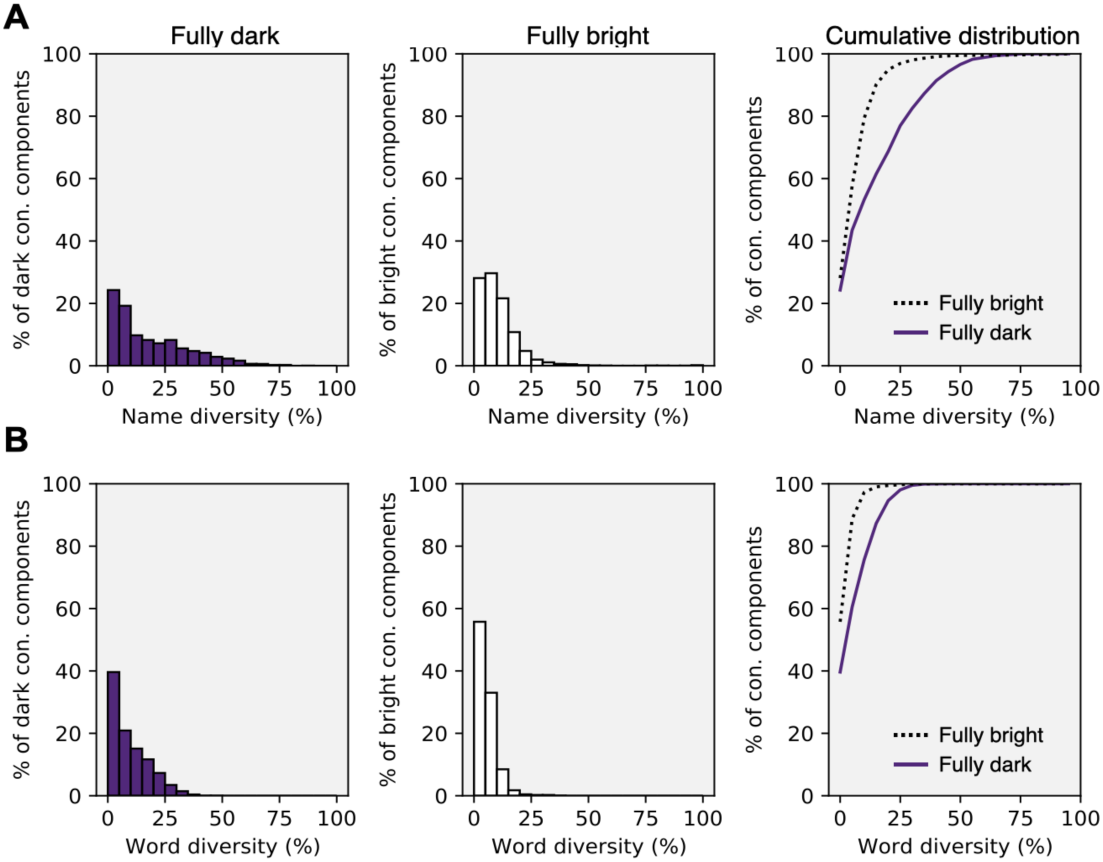
Diversity of the (A) names and (B) their word composition, for the individual fully dark and fully bright connected components. Name diversity is calculated as the number of unique protein names within a component by the total number of component proteins. Word diversity is calculated as the number of unique words across all protein names within a component by the total number of words, ignoring the words “protein”, “domain”, “family”, “containing”, and “superfamily”.

As expected, most bright connected components had a low semantic diversity, indicating a coherent and consistent naming. The maximum word diversity in bright connected components was 33%, which corresponds to cases where multiple varieties of the same name are present. One example is component 100’340, which is composed of multiple reviewed “Cytotoxin” names with different labels (e.g., Cytotoxin SP15a, Cytotoxin 10, etc.).

On the other hand, fully dark connected components tended to have a higher semantic diversity, with a name diversity of 19% (compared to 10% in fully bright components) and a word diversity of 7% (compared to 4% in fully bright components). The less diversely named dark components are those with previously submitted names not predicted by ProtNLM, such as component 159, where most proteins are named “DUF6516”. Conversely, the dark component with the highest semantic diversity was component 3’314, with a word diversity of 45%.

Component 3’314 is composed of 53 proteins that were given a wide variety of unrelated predicted names, from “Integrase” to “NADH-quinone oxidoreductase subunit F”, “Dynein light chain”, “Prophage protein”, etc. Despite their diverse predicted names, proteins in component 3’314 share a common fold (Fig. S3A) but find no structural homologs in the PDB. HMM searches over the PDB with HHpred highlighted a small local match to the tubulin-binding domain of *Chlamydomonas reinhardtii* TRAF3-interacting protein 1 (PDBID 5FMT, chain B) at a probability of 71%, but when clustered together, these two groups of proteins only form a few weak connections (Fig. S3A). We found a few members of component 3’314 dispersed throughout bacteria and bacteriophages, and they do not share a conserved genomic context (Fig. S3B). Together with the presence of prophage-associated protein encoding genes in these genomic contexts, such as “Host-nuclease inhibitor protein Gam” (Akroyd et al., 1986), these data support the “Prophage protein” title.

Our work highlights that the ProtNLM language model when presented with families with no homologs was hallucinating a diverse range of names. We identified 281 functionally dark components with a word diversity >20% and are now defining Pfam families for each of them (Mistry et al., 2020), with 22 already available in the upcoming Pfam release 36.0 (Table S1). This includes component 3’314, which gave rise to the PF21779 family and whose members are now titled DUF6874. As we define new Pfam families, the naming of these families should become consistent as future versions of ProtNLM consume this data. Starting from UniProt release 2023_01, the criteria for displaying ProtNLM names has changed to include an ensemble approach, an increased confidence threshold, and an automatic corroboration pipeline (https://www.uniprot.org/help/ProtNLM), thus many of these hallucinated names have reverted to “Uncharacterised protein”.

### 4. Structural outlier diversity in AFDB90

The availability of high quality predicted structures allows us to expand functional curation efforts to include structure-based similarities. One useful approach for this is structure-based alignment, which is described in (Barrio-Hernandez et al., 2023), and also incorporated into our examples to find similarities to proteins with known PDB structures using Foldseek. Here, we look into another angle of structure comparison, which is based on the concept of a “structural outlier”. By using an alphabet of substructure representations, covering 1’024 local structural contexts (16 residues in sequence and 10 Å spatial neighbourhood, Fig. S4), we trained an outlier detector on PDB structures and predicted the AFDB90 structures that have substructure compositions which are rare or absent in the PDB (See Methods Section 5). With this, 699’084 AFDB90 structures across 143’352 communities were predicted to be structural outliers, 30% of which are functionally dark (Fig. 7A). This analysis gives us a measure of plausibility that can help prioritise protein family definition. Importantly, in selecting the examples to explore in previous sections, we focused on structural inliers.

**Figure 7.**
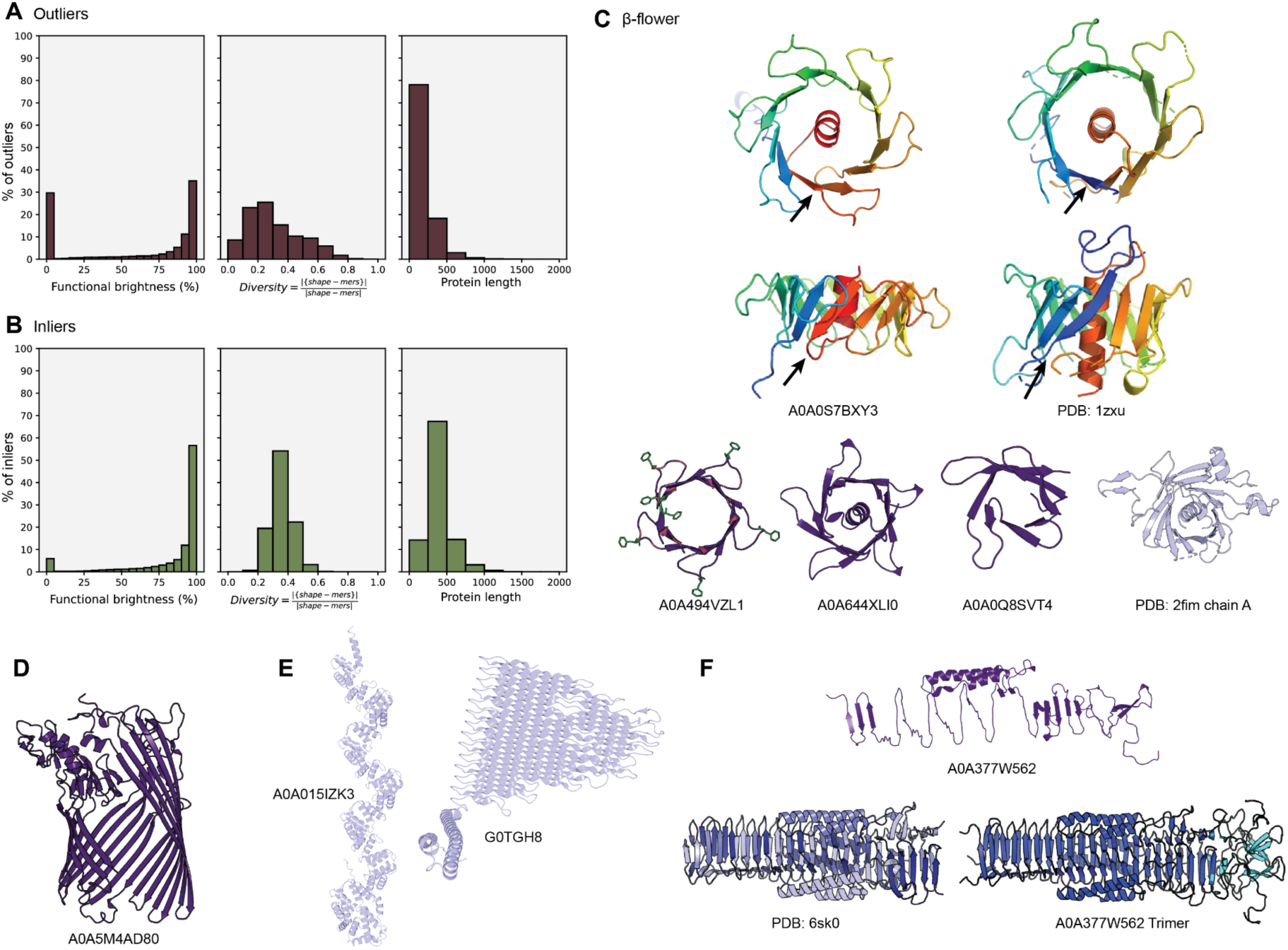
Structural outliers are diverse and can represent novel folds, protein fragments, repetitive proteins, or those that require folding conditions out of the scope of AlphaFold2. (A-B) Distribution of brightness, shape-mer diversity and length of the (A) structural outliers and (B) a set of structural inliers with the lowest outlier scores equivalent to the number of outliers. Shape-mer diversity is defined as the number of unique shape-mers by the length of the protein. (C) AlphaFold models of different variations of the β-flower, with positively charged residues in red and phenylalanine in green for A0A494VZL1, and PDB structures of *Arabidopsis thaliana* putative phospholipid scramblase (1zxu) and the human Tubby C-terminal domain (2fim). Black arrows indicate the circularly permuted loop in A0A0S7BXY3 and 1zxu (D) A structure outlier annotated as “TonB-dependent receptor-like” proteins that is a fragment of a β-barrel domain. (E) Two long repetitive outliers, one belonging to the Tetratricopeptide-like helical domain superfamily (A0A015IZK3) and one to the PE-PGRS superfamily (G0TGH8). (F) AlphaFold model of an outlier annotated as containing “Putative type VI secretion system, Rhs element associated Vgr domain” (A0A377W562), a trimeric PDB structure (6sk0) also containing this domain, and an AlphaFold-Multimer model of the A0A377W562 trimer.

We observed that structural outliers tend to be shorter and more repetitive than inliers (Fig. 7A,B), indicating that such an approach could be applied to predicted structures to annotate fragments and low complexity proteins, some of which could contain sequencing errors or not have translation-level evidence. Outliers can also represent novel folds, as in the example below.

### 4.1. The β-flower fold

UniRef50_A0A494VZL1 is an example of a structural outlier without homologs in the PDB and whose AFDBv4 homologs are found only at the structure level. A0A494VZL1 folds as a shallow, symmetric β- barrel domain with 96 residues. This barrel is formed by 10 short antiparallel β-strands that form a hydrophobic channel. On one side of the β-barrel, the loops connecting each strand are much longer (9 residues) than those on the other side (4 residues), which make it look like a flower (Fig. 7C). For this reason, we named it “β-flower”. Some of these loops are enriched with positively charged arginine and lysine residues, and at the tip contain phenylalanines whose side-chains point towards the exterior of the β-barrel.

While UniRef50_A0A494VZL1 is a singleton in our network (i.e., it does not belong to any connected component with at least 2 nodes), a Foldseek search on structural representatives of AFDB90 (the AFDB90Communities set, defined in the *Material and Methods* section) found hits with a TM-score better than 0.6 to 43 UniRef50 clusters in AFDB90. 39 of these are from bacteria, and map into 13 different sequence-based connected components with 1 to 6 nodes, 39 are structural outliers and 13 are bright because they are annotated almost full-length as “Cell wall-binding protein” or “MORN repeat variant”. Within these 43 UniRef50 clusters, there are at least three globally different folds, which are variations of that of UniRef50_A0A494VZL1 (Fig. 7C). These differ particularly on the number of strands (8, 10, or 12) and, consequently, the number of long loops (4, 5 or 6), which makes these domains look like square, pentagonal or hexagonal “flowers”. The “petals” of these flowers comprise beta-hairpins that are arranged in four-, five- or six-fold symmetry. However, within the structural homologs there are also some that resemble half of a flower, perhaps corresponding to fragments of longer domains.

Many of the proteins in this family possess N-terminal lipoprotein attachment motifs (Hayashi and Wu, 1990; Klein et al., 1988), suggesting they are associated with the bacterial inner membrane or transferred to the inner leaflet of the outer membrane. Although no structural similarity to the PDB was highlighted by Foldseek and HMM-HMM searches with HHpred also returned no results, the β-flower folds with six-fold symmetry are reminiscent of the Tubby C-terminal domain (Bateman et al., 2009), that adopts a twelve-stranded β-barrel fold enclosing a central hydrophobic helix (Fig. 7C). Besides the global structural similarity, these proteins share a common network of aromatic hydrophobic residues that flank the edges of the β-strands and point toward the interior of the β-barrel, thus engaging in tight contacts with the central hydrophobic helix. Interestingly, this feature is also preserved in the proteins lacking the central helix in the barrel interior, e.g. A0A494VZL1.

There are, however, some topological differences that are more evident when comparing the β-flowers with 6-fold symmetry to the Tubby-like domains (Fig. 7C). The N-terminal strand of Tubby domains is circularly permuted in the β-flowers, where its position is occupied by the C-terminal strand. The presence/absence of the C-terminal strand in the β-barrel leads to a different entry point of the C-terminal hydrophobic helix into the channel formed by the β-barrel, which defines the difference in its directionality. Other differences pertain to the length of the β-strands and the connecting loops on one side of the β-barrel, which in the β-flower proteins are significantly shorter. Tubby-like proteins either bind to phosphoinositides or function as phospholipid scramblases (Bateman et al., 2009) and this provides a functional hypothesis to test for these proteins. The diversity of these proteins has been added to Pfam as the new entries PF21784, PF21785 and PF21786, which together with the Tubby C-terminal domain now form the CL0396 clan.

### 4.2. Fragments, repeats and obligate complexes

Structural outliers can also represent fragments of existing families (Fig. 7D), highly repetitive proteins that are rare in the PDB (Fig. 7E), and proteins which require conditions to fold, such as binding partners, not modelled by AlphaFold2 (Fig. 7F).

9’519 AFDB90 structural outliers, across 1’258 components mapping to 16’524 AFDBv4 proteins, are annotated by InterPro as containing the ‘TonB-dependent receptor-like’ (TBDT) domain, of which 86% are fully bright. TBDTs have 22-stranded β-barrel C-terminal domain (Noinaj et al., 2010) but most of these outliers have incomplete β-barrel domains (e.g. Fig. 7D), with 82% having less than the required number of β-sheet shape-mers found in the 45 PDB structures from (Noinaj et al., 2010), despite 55% not being explicitly annotated as fragments in UniProtKB. These could be due to frameshift errors introduced in whole-genome sequencing runs. Many of these have significant Foldseek hits to the PDB with TM-score > 0.7, indicating that relying only on automated similarity scores from sequence and structure alignment may still lead us astray for functional annotation.

Long repetitive proteins are also marked as outliers, as they are rare or absent in the PDB (FIG. 7E). 6’791 outliers have over 500 residues and a shape-mer diversity fraction < 0.1 of which 4’948 are fully bright. Many of these belong to the ‘Tetratricopeptide-like helical domain superfamily’ where the median PDB structure length of structures with resolution < 3Å is only 370. Another example of repetitive structural outliers which are thought to be novel folds include the unusual “PGRS domains”, found widely in mycobacteria (Berisio and Delogu, 2022).

Thirty-six AFDB90 structural outliers across 2 components, mapping to 103 AFDB proteins, are labelled as having the “Putative type VI secretion system, Rhs element associated Vgr domain”, for which only 3 Cryo-EM PDB structures are known. The models do not resemble the PDB structures (Fig. 7F) because these proteins are obligate complexes that form trimeric β-solenoids and thus the predicted fold of the monomeric chains, which is what AFDB provides, is not functionally meaningful, while the AlphaFold-Multimer model of the trimer (Fig 7F) has 1.1Å RMSD to the PDB structure.

Although structural information in AFDB is highly informative and novelties in terms of protein families can be detected, our results highlight that the protein structure space as predicted by AlphaFold needs to be put into evolutionary, functional, and structural context before any model is used as a reference structure, even if they have high predicted accuracy.

## Conclusions: Towards large-scale protein function annotation

In this work, we carried out a large-scale analysis of the catalogued protein sequence space covered by predicted high quality structural information, as made available through AlphaFold database version Our results indicate that 19% of this space is composed of protein families and superfamilies that are functionally dark and can not, solely on the basis of sequence similarity, be annotated to a known protein family. We also highlight that a large fraction of these can not be named consistently using the most recent deep-learning-based approaches and demonstrate that functional annotation of proteins, even from a purely computational perspective, requires a combination of data sources and approaches.

Our findings highlight the power of large-scale similarity networks to accurately annotate protein function. In cases where simple homology transfer methods are not effective, pooling information from across the network can enhance remote homology detection. When combined with traditional protein evolution approaches, genomic context information, structural outlier measures, structure-based function prediction and predicted name diversity, we can gain valuable insights into putative protein function, annotate undescribed protein families that can be prioritised for experimental characterization, and identify false annotations or incorrect protein sequences. However, it is crucial to combine individual predictions from these tools with statistics of predictions across similar proteins to increase the confidence of the annotation.

It is important to mention, however, that our study has some caveats and limitations. Firstly, all the analyses presented, including the sequence-based clustering and outlier analysis, depend on thresholds and parameters that were chosen based on prior experience and not optimised for any particular downstream application. All alignment similarity techniques used considered coverage across the full protein sequence, in order to enable detection of novel full-length protein families. This indicates that some remote homologies may have been missed, with a domain-based exploration providing a possible complementary solution. In addition, our highly particular definition of functional brightness excluded predicted intrinsically disordered and coiled-coil proteins, and misclassifies some functionally uncharacterised proteins as “bright” because of the presence of ambiguous annotations such as “transmembrane” or “repeat” domains or some characterised ones as “dark” because their annotations are “Putative” or they were not assigned a family. Thus, the examples we discuss are the low-hanging fruit of uncharacterised or unannotated protein families, but they are only the tip of the iceberg. Still, we have found that these clusters are a rich source of new Pfam families, and have begun to add a selection of them to the upcoming release 36.0.

Recently, several complementary approaches have been developed to categorise the diversity of the protein universe and uncover novelties (Akdel et al., 2022; Barrio-Hernandez et al., 2023; Bordin et al., 2023), again highlighting the importance of incorporating multiple perspectives and methods in protein function annotation. These approaches showcase the significance of using a diverse set of information to gain a more complete understanding of protein function and its role in cellular processes. We expect that further advances in deep-learning-based methods for function prediction (Gligorijević et al., 2021), remote homology detection (Kaminski et al., n.d.; Pantolini et al., 2022) and large-scale protein structure prediction (Lin et al., n.d.) will allow in the near future to take these analyses to a much larger scale and at a much finer resolution.

## Acknowledgements

We would like to thank the SWISS-MODEL development team for technical support and text revisions, Gemma C. Atkinson, Tomasz Kościółek, Lydie Lane, Max Bileschi, and Lynne Regan for insightful discussions and comments, the Cosmograph team for providing the fastest network graph visualization tool that works in the browser, and sciCORE at the University of Basel for providing computational resources and system administration support.

This work was supported by funding from the SIB - Swiss Institute of Bioinformatics (https://www.sib.swiss/), the Biozentrum of the University of Basel (https://www.biozentrum.unibas.ch/), by the European Union via project MIBEst H2020-WIDESPREAD-2018-2020/GA number 857518 (T.T. and V.H.), by a grant from the Estonian Research Council (PRG335 to T.T. and V.H.), the Knut and Alice Wallenberg Foundation (2020-0037 to V.H.), Swedish Research Council (Vetenskapsrådet) grants (2021-01146 to V.H.), Cancerfonden (20 0872 Pj to V.H.), and the Biotechnology and Biological Sciences Research Council and the NSF Directorate for Biological Sciences (BB/X012492/1 to A.B).

## Materials and Methods

### 1. Data collection

We started from the 53’625’855 UniRef50 (Suzek et al., 2015) clusters as of August 2022 (UniRef version 2022_03) and the 214’683’829 structural models for most UniProtKB entries available via the AlphaFold database (version 4, AFDBv4). For each Swiss-Prot (Boutet et al., 2016), TrEMBL (UniProt Consortium, 2023) and UniParc (Leinonen et al., 2004) entry in each UniRef50 cluster we collected their sequence, taxonomy and functional and structural annotations from UniProt and InterPro (Paysan-Lafosse et al., 2023). This includes domains, protein families, predicted transmembrane segments, predicted signal peptides, and predicted disorder and coiled coil regions. Redundant, overlapping domain, families, coiled coil and disorder annotations were continuously merged (Fig. 1A), selecting as the preferential name the first occurrence that did not include “Putative”, “Hypothetical”, “Uncharacterised” and “DUF”. In parallel, we also mapped each entry in AFDBv4 to their corresponding UniRef50 cluster, selecting as the structural representative of that cluster the longest protein with an average pLDDT better than 70.

### 2. Darkness estimation

We define functional brightness of a given protein as the full-length coverage with annotations of its close homologs, with 0% meaning “dark” and 100% meaning “bright”. For that, we first computed the full-length coverage with annotations for all entries in all UniRef50 clusters, and then considered a UniRef50 cluster as “bright” as the “brightest” sequence it encompasses (Fig. 1A). Annotations considered were: domains as annotated in InterPro, and families, predicted disorder and predicted coiled coil regions as annotated in UniProtKB and UniParc. For any domains and families, all those with “Putative”, “Hypothetical”, “Uncharacterised” and “DUF” in their name were given a coverage of zero and, thus, considered “dark”.

### 3. Large-scale sequence similarity network

To model the sequence landscape covered by all UniRef50 clusters with a high-confidence structural model, we built a large-scale sequence similarity network of 6’136’321 UniRef50 clusters with a structural representative with a pLDDT (Jumper et al., 2021) better than 90 (AFDB90 dataset). For that, all-against-all MMseqs2 (Steinegger and Söding, 2017) comparisons were carried out with the sequence representatives of all selected UniRef50 clusters, linking two sequences if they find a match that covers at least 50% of their full length sequences with an E-value better than 10^−4^. Each edge was given a weight proportional to the E-value of the match and, to facilitate the handling of the resulting network, a maximum of 4 outbound edges were considered per node. For that, nodes were sorted based on their label (the UniProtID of the corresponding protein) so that edges always adopt a downward direction in the list (e.g., A→B and B→A always correspond to A→B) and for each node only the 4 outbound higher weight edges considered (Fig. 1B). The direction of the edges was not considered for further analysis.

To further visualise the graph, each connected component in the graph was simplified to a set of connected communities. For that, per connected component, communities of highly connected nodes were detected using the asynchronous label propagation algorithm, as implemented in the *asyn_lpa_communities* method in networkx (Hagberg et al., 2008). This reduced the graph to a total of 688’852 communities (hereafter referred to as the AFDB90Communities set) connected by a total of 1’488’764 edges, whose layout could then be computed with Cosmograph (https://cosmograph.app/). For the simulation with Cosmograph, the maximum space allowed (8192), a gravity of 0.5, a repulsion of 1.4, a repulsion theta of 1.71, a link strength of 2, a minimum link distance of 1 and friction of 1 were used. For each community, we then collected the longest and median-length representatives, whose structures were later used in section 5. For visualisation, individual connected components were extracted from the layouted graph and drawn with Datashader (https://datashader.org/index.html). The interactive web version of this network was created using the Cosmograph library for network visualisation and the Mol* toolkit for 3D macromolecular visualisation of individual structure representatives.

### 4. Sequence-based prioritisation of dark connected components and their semantic name diversity

Each node in a connected component in our non-simplified network was attributed a functional brightness value, and for each connected component the average brightness was computed. Connected components were sorted by their average brightness and their overall size (i.e., number of nodes), so that the top ranking were the largest, darkest connected components. We selected all connected components with more than 1 node and a median brightness below 5%.

As during the time this work was being carried out UniProt implemented a deep learning-based approach for naming all those proteins from TrEMBL titled as “Uncharacterised protein” (ProtNLM) (Gane et al., 2022), we analysed the consistency of the names predicted for the proteins included in each of the selected dark connected components. For this, we did not consider only the individual UniRef50 representatives, but all unique sequences included in all those clusters that make the connected component, and considered only those connected components that account for more than 50 unique protein sequences. For each fully dark (functional brightness ≤ 5%) and fully bright (functional brightness ≥ 95%) component, we collected the name of the representative of each UniRef100 cluster included as of UniProt version 2022_04 (December 2022), and computed the proportion of unique names (i.e., name diversity) as well as the proportion of unique words (i.e., word diversity), in order to account for small variation of the same name. By comparing the distribution of name and word diversities between bright and dark components, this allowed the identification of dark proteins that are similar at the sequence level but not named consistently, likely representing protein families that ProtNLM has never seen before and thus unable to name.

### 5. Structural and substructural features of dark proteins and novel protein families

To represent and analyse 3D substructure composition, we build upon Geometricus (Durairaj et al., n.d.), and use 16 rotation invariant moments from (Flusser et al., 2016, 2003; Mamistvalov, 1998) and the chiral invariant moment from (Hattne and Lamzin, 2011). These moments were calculated for 4 different fragmentation types on ɑ-carbon coordinates: *k*-mers of size 8 and 16, and spheres of radii 5Å and 10Å; for a total of 68 moments for each central residue in a protein. We then trained a neural network using PyTorch (Paszke et al., 2019) with these 68 moments as input, 2 linear hidden layers of size 32 and a sigmoid output layer of size 10, and with contrastive loss to reduce the output distance between equivalent pairs of central residues and increase the distance between non-equivalent pairs in a training set. As the output of the network is 10 floating point numbers between 0 and 1, this could be discretized into 10 bits based on whether the value was greater than or less than 0.5, resulting in 1024 shape-mers.

The training set is created from structures from the CATH database having less than 40% sequence identity (CATH40) that could be assigned to a CATH functional family (FunFam) with an E-value < 1×10^−6^. From these 8’333 structures, US-align (Zhang et al., 2022) was used to align and superpose all pairs within each FunFam cluster and 3×8’333 randomly chosen pairs across clusters. Aligned pairs of residues from two proteins belonging to the same FunFam with an alignment TM-score > 0.8 were considered as positive pairs. Aligned or random pairs of residues from two proteins belonging to different CATH superfamilies, with an alignment TM-score <0.6 were considered as negative pairs. In addition, using all 31,883 CATH40 proteins, we sampled up to 50 pairs of central residues from each protein, where positive pairs had <2 sequence distance and negative pairs had 5-20 sequence distance. In total, this resulted in 6 million residue pairs for training, of which 42% were positive pairs. This dataset could be used for training and/or refining any kind of residue-level contrastive learning task. Training took 30 mins on 1 RTX-3080TI with the ADAM optimizer, a batch size of 1024, and a learning rate of 10^−3^ over 5 epochs.

Shape-mers were calculated for all AFDB90 proteins, and all ProteinNet CASP12 proteins in the 100% sequence identity set (AlQuraishi, 2019). For AlphaFold models, we followed the approach described in the analysis of AFDBv1 (Akdel et al., 2022) to split each protein into segments with Gaussian smoothed plDDT > 70, after first splitting into domains based on a combination of pLDDT and the predicted aligned error (PAE) matrix. Shape-mers were then calculated for each segment in each domain and concatenated. Proteins with less than 20 amino acids were ignored. A shape-mer diversity fraction was defined for each protein as the number of unique shape-mers divided by the total number of shape-mers (which is equal to the protein length if all residues have high pLDDT). Fig. S4 shows an example AlphaFold protein with its 6 most common shape-mers highlighted.

We trained a FastText model (Bojanowski et al., 2017) on the shape-mer bit representations from the ProteinNet CASP12 dataset using Gensim (Rehurek and Sojka, n.d.)(v4.2.0) with a window size of 16 and embedding size of 1024. Fig. S5A shows the sensitivity of SCOPe family retrieval on the SCOPe40 dataset of 11’211 structures for all-vs-all Smith-Waterman alignment with FastText shapemer similarities used as the score matrix (runtime: 12 mins on 10 threads). Shape-mer FastText alignment scores are compared to three structure aligners, DALI (Holm, 2020), Foldseek (van Kempen et al., n.d.), and TM-align (Zhang and Skolnick, 2005); one sequence aligner, MMseqs2 (Steinegger and Söding, 2017); and 2 other structure alphabet-based structural sequence aligners, 3D-BLAST (Mavridis and Ritchie, 2010) and CLE-SW (Wang and Zheng, 2008), using the scripts and benchmark data provided in (van Kempen et al., n.d.). The benchmark results demonstrate that the learned structural alphabet and FastText similarities still have discriminative power in distinguishing protein families, despite being much less “local” than approaches such as Foldseek and TM-align which work on individual coordinates of up to 2 residues. We don’t explore further alignment optimization, such as compositional bias correction or penalty optimization to increase sensitivity, as more local structural aligners will still have the advantage of higher resolution alignment. However, for the task at hand, our substructure representations give us a good compromise - a discriminative structural alphabet for representing a protein structure as a structural sequence; and substructure decomposition at the level of whole secondary-structural elements, allowing for a broader exploration of substructure composition across the AlphaFold database.

To compare substructure compositions, we obtained protein-level embeddings by averaging across normalised FastText embeddings using the *get_sentence_vector* function. Fig. S5B shows the distributions of cosine distances of these embeddings within the same SCOPe family and across SCOPe folds, demonstrating that they have discriminative power in representing substructure compositions. We trained the Isolation Forest algorithm (Liu et al., 2008) as implemented in scikit-learn v1.1.1 (Pedregosa et al., 2011) on the ProteinNet CASP12 FastText sentence embeddings with 1% contamination rate, and used this trained model to predict structural outlier scores for proteins in the AFDB90 dataset. Proteins with a negative score are labelled as outliers.

### 6. Computational investigation of selected examples

We further investigated 4 different types of examples of dark proteins and connected components we encountered in our dataset: (1) a dark connected component whose models are structural inliers and are named consistently, but have no matches in the PDB (component 159), (2) a dark component whose models are structural inliers, are named consistently and have structural homologs in the PDB (component 27), (3) a dark connected component that is named inconsistently (component 3314), and (4) examples of different kinds of structural outliers, including a novel fold (β-flower fold). In all cases, we combined data from the sequence-based network and its functional brightness annotations, as well as from structural searches with Foldseek and the outlier scores. Structural homologs for selected representatives (those with a length close to the median length in the component) in the PDB or the AFDB90Communities set were searched with Foldseek using the TM-align mode (van Kempen et al., n.d.). Remote sequence homologs were detected for selected representatives by HHPred searches over the PDB, ECOD and Pfam databases through the MPI Bioinformatics toolkit using default settings (Gabler et al., 2020; Pereira and Alva, 2021). Further case-by-case analyses were also carried out as described below.

### 6.1. Connected component 159

Ninety-four randomly selected sequences from component 159 were aligned with MUSCLE (Edgar, 2004). The resulting alignment was used for three independent PSI-BLAST (Altschul et al., 1997) searches over the eukaryotic, archaea and bacterial sequences in *nr* (nr_euk, nr_arc, nr_bac) with 8 rounds through the MPI-Bioinformatics toolkit as of October 2022 (Gabler et al., 2020; Pereira and Alva, 2021). All collected sequences were filtered to a maximum sequence identity of 95% with MMseqs2 (Steinegger and Söding, 2017) and clustered based on BLASTp all-against-all pairwise searches with CLANS until equilibrium at an E-value of 1×10^−10^.

The resulting sequence similarity network was used as input for GCsnap (v1.0.17) (Pereira, 2021) for the analysis of the conservation of the genomic contexts encoding for each of the proteins in the individual clusters. A window of four flanking genes was used, MMseqs2 was employed for protein family clustering at an E-value better than 1×10^−4^ and clusters of similar genomic contexts were detected using the *operon_cluster_advanced* method, which employs PaCMAP (Wang et al., 2020) to project genomic contexts in 2D based on their family composition and DBSCAN (Ester et al., 1996) (as implemented in scikit-learn) to identify clusters of similar genomic contexts. For this, only families that were found in at least 30% of all genomic contexts were considered. For each cluster in the sequence similarity network and each identified neighbour family, up to 100 structure representatives were selected from AFDB and used as input for DeepFRI with default settings (Gligorijević et al., 2021). The top 10 most common predictions per cluster/context family were retrieved. The highest average scoring and most frequent predicted for each case were considered the most likely molecular functions for each case.

We further predicted the 3D structure of the complex between a representative of the main cluster in the network (subcluster 1a), which we selected for further experimental validation and characterisation as described below. For that, AlphaFold-Multimer (Evans et al., 2021) version 3 was used to generate 3D models of a tetramer consisting of two chains of the *Allochromatium tepidum* TumE toxin (EntrezID: WP_213381069.1) and two of its putative, cognate TumA antitoxin (EntrezID: WP_213381068.1) with default settings and relaxation. The model with the best predicted TM score (pTM) and interface pTM score was selected (Evans et al., 2021).

### 6.2. Connected component 27

All non-redundant protein sequences represented by the nodes of connected component 27 (i.e., all UniRef100 representatives) were collected and filtered to a maximum sequence identity of 50% with MMseqs2. The reduced set of sequences was aligned with MUSCLE and the resulting MSA was used as input for three independent BLAST searches over the eukaryotic, archaea and bacterial sequences in *nr* filtered to 70% sequence identity (nr_euk70, nr_arc70, nr_bac70) through the MPI-Bioinformatics toolkit as of January 2023. The same BLAST searches were carried out for SWISS-PROT representatives of the PglB, STT3 and YfhO families (UniProtIDs PGLB_CAMJR, STT3_YEAST and YFHO_BACSU). The full-length sequences matched in all searches were then combined with those representatives of connected component 27 and filtered to a maximum sequence identity of 30% with MMseqs2. The resulting set of 7’004 sequences was clustered based on BLASTp all-against-all searches with CLANS at an E-value of 1×10^−20^ until equilibrium.

### 6.3. Connected component 3314

All non-redundant protein sequences represented by the nodes of connected component 27 (i.e., all UniRef100 representatives) were collected and filtered to a maximum sequence identity of 50% with MMseqs2. The reduced set of sequences was aligned with MUSCLE and the resulting MSA was used as input for four independent BLAST searches over the eukaryotic, archaea, bacterial and viral sequences in *nr* filtered to 70% sequence identity (nr_euk90, nr_arc90, nr_bac90, nr_vir90) through the MPI-Bioinformatics toolkit as of January 2023. The same BLAST searches were carried out for the tubulin-binding domain of *Chlamydomonas reinhardtii* TRAF3-interacting protein 1 (UniProtID A8JBY2_CHLRE, residues 1-131). The full-length sequences matched to component 3314 homologs and the local sequence matched to TRAF3-interacting protein 1 tubulin binding domain were then combined with those representatives of component 3314 and filtered to a maximum sequence identity of 90% with MMseqs2. The resulting set of 890 sequences was clustered based on BLASTp all-against-all searches with CLANS at an E-value of 1×10^−5^ until equilibrium. The 141 sequences making subcluster 1 in the resulting network, which included the component 3314-like proteins, were extracted from the network, filtered to a maximum sequence identity of 50% with MMseqs2 and used as input for GCsnap (v1.0.17), where a window of four flanking genes was used and MMseqs2 employed for protein family clustering at an E-value better than 1×10^−4^.

### 6.4. β-flower fold

We constructed three new Pfam families to cover the sequence space of β-flower proteins. To do this we selected example proteins with 4,5 and 6-fold rotational symmetry and iteratively searched for homologs using the hmmsearch software of the HMMER package (version 3.3) (Eddy, 2011). In general, we used an inclusion threshold of 27 bits, but manually lowered the threshold to identify more homologs or raised it to exclude false matches as identified by AlphaFold2 models. These three families were added to Pfam with accession numbers: PF21784, PF21785 and PF21786 and these families were added to Pfam clan CL0396, which includes the Tubby C-terminal domain. The families were included in the Tubby C-terminal clan considering their significant structural similarity to known structures of Tubby-like proteins, in particular the structure of plant Tubby-like At5g01750 protein (PDBID 1zxu).

### 7. Experimental validation and characterisation of a predicted toxin-antitoxin family (component 159)

Six Proteobacteria TumE examples from subcluster 1a in the CLANS sequence similarity network produced in 6.1. and their cognate TumA antitoxins were selected for experimental characterization (Supplementary file 2). The plasmids were constructed using the Circular Polymerase Extension Cloning (CPEC) (Quan and Tian, 2011) approach with synthetic DNA procured from Integrated DNA Technologies. ORFs were synthesised with added strong Shine-Dalgarno sequence (AGGAGGAATTAA) and flanking sequences overlapping with multicloning sites of pBAD33 (toxin genes) or pMG25 (antitoxin genes). The DNA fragments were amplified with Phusion polymerase (Thermo Scientific™) using pBAD_SD_TOX_fwd and pBAD_TOX_MCS_rev or pMG25_insert_fwd and pMG25_insert_rev primer pairs. pBAD33 was linearized using primers pBAD_lin_1 and pBAD_lin_2 and pMG25 was linearized using pMG25_lin_from_BlpI and pMG25_lin_from_HindIII. CPEC with Phusion polymerase (Thermo Scientific™) was performed to clone the genes into the vector backbone (25 cycles with 5 min 30 s extension). The CPEC reaction mixture was transformed into DH5α *E. coli* cells and colony PCR with HOT FIREPol® Blend Master Mix (Solis Biodyne) was used to identify colonies with correctly sized inserts. Plasmids were extracted from the overnight cultures using FavorPrepTM Plasmid Extraction Mini Kit (Favorgen) and sequenced. The cognate antitoxin plasmid or empty pMG25 was co-transformed with the toxin plasmids into BW25113 *E. coli* cells. DNA fragments and DNA oligonucleotides used for plasmid construction are provided in Supplementary file 2.

Validation of toxicity and metabolic labelling experiments with ^35^S methionine, ^3^H uridine and ^3^H thymidine were performed as described earlier (Kurata et al., 2022). Briefly, *E. coli* BW25113 strains were transformed with a plasmid pair that allowed for controllable co-expression of putative TumE toxins (pBAD33 derivatives, the toxin is expressed under the control of L-arabinose-inducible P_BAD_ promotor) and TumA antitoxins (pMG25 derivatives (Jaskólska and Gerdes, 2015), IPTG-inducible expression of the antitoxin is driven by P_Tac_ promotor) and pregrown in liquid Lysogeny broth (LB) medium (Lennox) supplemented with 100 µg/mL carbenicillin (AppliChem) and 25 µg/mL chloramphenicol (AppliChem) as well as 0.2% glucose (for repression of toxin expression). Serial 10-fold 5 µL dilutions were spotted on LB plates supplemented with antibiotics (carbenicillin and chloramphenicol) as well as either 0.2% glucose (repressive conditions) or 0.2% arabinose and 1 mM IPTG (induction conditions). Plates were scored after an overnight incubation at 37 °C.

For metabolic labelling experiments with TumE toxins, *E. coli* BW25113 strains co-transformed with pBAD33 derivatives (for L-arabinose-inducible expression of toxins) as well as the empty pMG25 vector were first plated out on LB plates supplemented with 100 µg/ml carbenicillin, 25 µg/ml chloramphenicol and 0.2% glucose (to suppress the leaky expression of the toxin). Using fresh, individual *E. coli* colonies for inoculation, 2 mL liquid cultures were prepared in defined Neidhardt MOPS minimal media (Neidhardt et al., 1974) supplemented with 100 µg/ml carbenicillin, 25 µg/ml chloramphenicol, 0.1% of casamino acids, and 0.2% glucose, and grown overnight at 37 °C with shaking. Next, experimental 15-mL cultures were prepared in 125 mL conical flasks in MOPS medium supplemented with 0.5% glycerol, 100 µg/ml carbenicillin, 25 µg/ml chloramphenicol as well as a set of 19 amino acids (lacking methionine), each at final concentration of 25 µg/mL. These cultures were inoculated overnight to final OD_600_ of 0.05, and grown at 37 °C with shaking up to of OD_600_ 0.2. At this point, one 1-mL aliquot (the pre-induction zero time-point) was transferred to 1.5 mL Eppendorf tubes containing 10 µL of radioisotope – either ^35^S methionine (4.35 µCi, Perkin Elmer), or ^3^H uridine (0.65 µCi, Perkin Elmer) or ^3^H thymidine (2 µCi, Perkin Elmer) – and transferred to the heat block at 37 °C. Immediately after, the expression of toxins in the remaining 14 mL culture was induced by addition of L-arabinose (final concentration of 0.2%). Throughout the toxin induction time course, 1-mL aliquots were taken from the 15 mL culture and transferred to 1.5 mL Eppendorf tubes containing 10 µl of radioisotope (^35^S methionine / ^3^H uridine / ^3^H thymidine). The incorporation of radioisotopes was stopped after 8 minutes of incubation at 37 °C by adding 200 µL of ice-cold 50% trichloroacetic acid (TCA) to 1 mL cultures. In parallel with taking the time-points for labelling, 1 mL aliquots were taken for OD_600_ measurements. Isotope incorporation was quantified by normalising radioactivity counts (CPM) to OD_600_, with the pre-induction zero time-point set as 100%. All experiments were performed in three biological replicates (i.e. using three independent cultures inoculated from three different colonies).

## Data and code availability

All the code to collect and process the annotation data in UniProtKB, UniParc and InterPro, and the pLDDT data from AFDB is available at https://github.com/ProteinUniverseAtlas/dbuilder. All analysis code, data and metadata generated for the current submission is available at https://github.com/ProteinUniverseAtlas/AFDB90v4. In addition, an interactive version of the sequence similarity network, queriable by UniProt ID, protein title, connected component and community ID, is available at https://uniprot3d.org/atlas/AFDB90v4.

## SUPPLEMENTARY FIGURES

**Figure S1.**
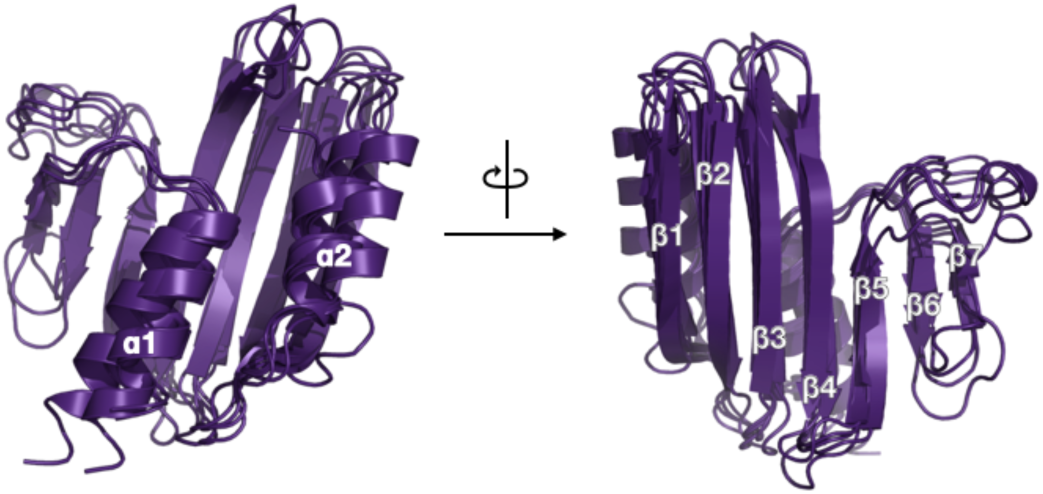
Structural conservation and structure-based function prediction of TumE. Structural superposition of five randomly selected members of component 159 (UniProtIDs A0A0E3S9F7, A0A3R7AQ40, A0A520JWH3, A0A1W9UY89, A0A7J4P9B0) with secondary structure elements labelled.

**Figure S2.**
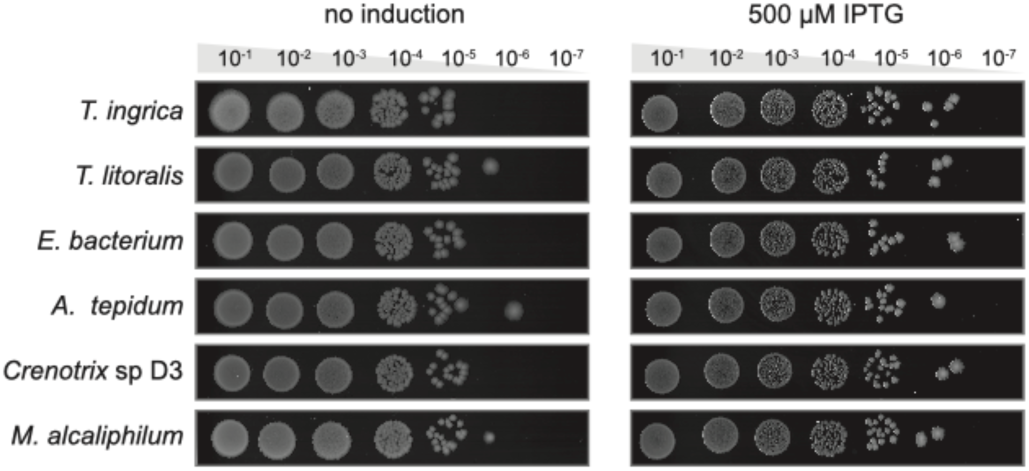
Testing the toxicity of putative TumA antitoxins. Antitoxin expression plasmids were cotransformed with empty toxin expression vectors (pBAD33) into *E. coli* BW25113 cells. The bacteria were grown for five hours in liquid LB media supplemented with appropriate antibiotics. The cultures were normalized to OD600 = 1.0, serially diluted and spotted on LB agar plates containing appropriate antibiotics and 500 µM IPTG for antitoxin induction and 0.2% arabinose to mimic the conditions in toxin neutralisation assay. The experiment was made in three biological replicates.

**Figure S3.**
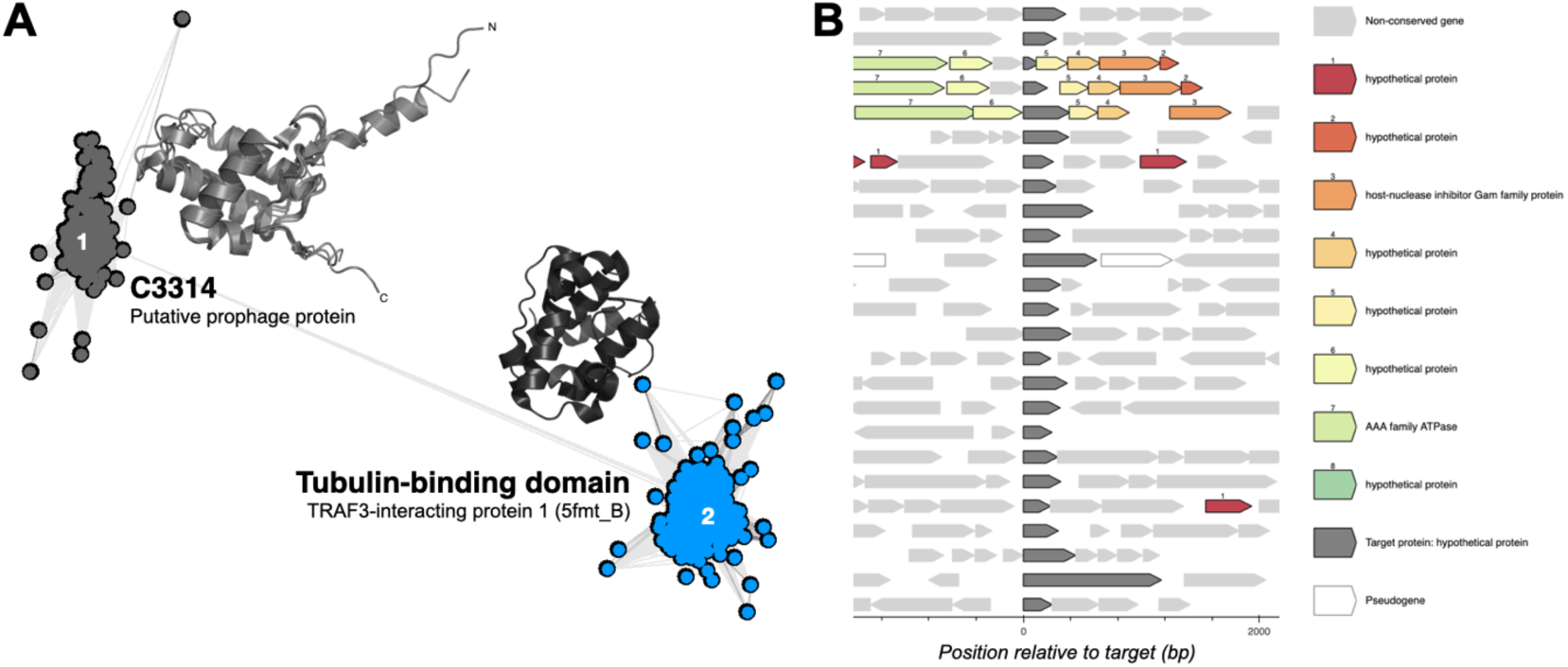
The highly semantically diverse prophage-associated connected component 3314. (A) Sequence similarity network of homologs of members of connected component 3314 and the tubulin-binding domain of TRAF3-interacting protein 1, as computed with CLANS at an E-value threshold of 1×10^−5^. Points represent individual proteins and grey lines BLASTp matches at an E-value better than 1×10^−4^. Individual subclusters are labelled 1-2 and structural representatives are shown. For subcluster 1, 5 randomly selected structural representatives of component 3314 are superposed (UniProtIDs A0A0F9A5W1, A0A0P9GTS8, AOA418VYX3, A0A2S5M855, A0A2K2VML8). For subcluster 2, the tubulin-binding domain of *Chlamydomonas reinhardtii* TRAF3-interacting protein 1 (PDBID 5fmt, chain B) is shown. (B) Genomic context conservation of 30 sequences from subcluster 1 with a maximum sequence identity of 30%, as computed with GCsnap.

**Figure S4.**
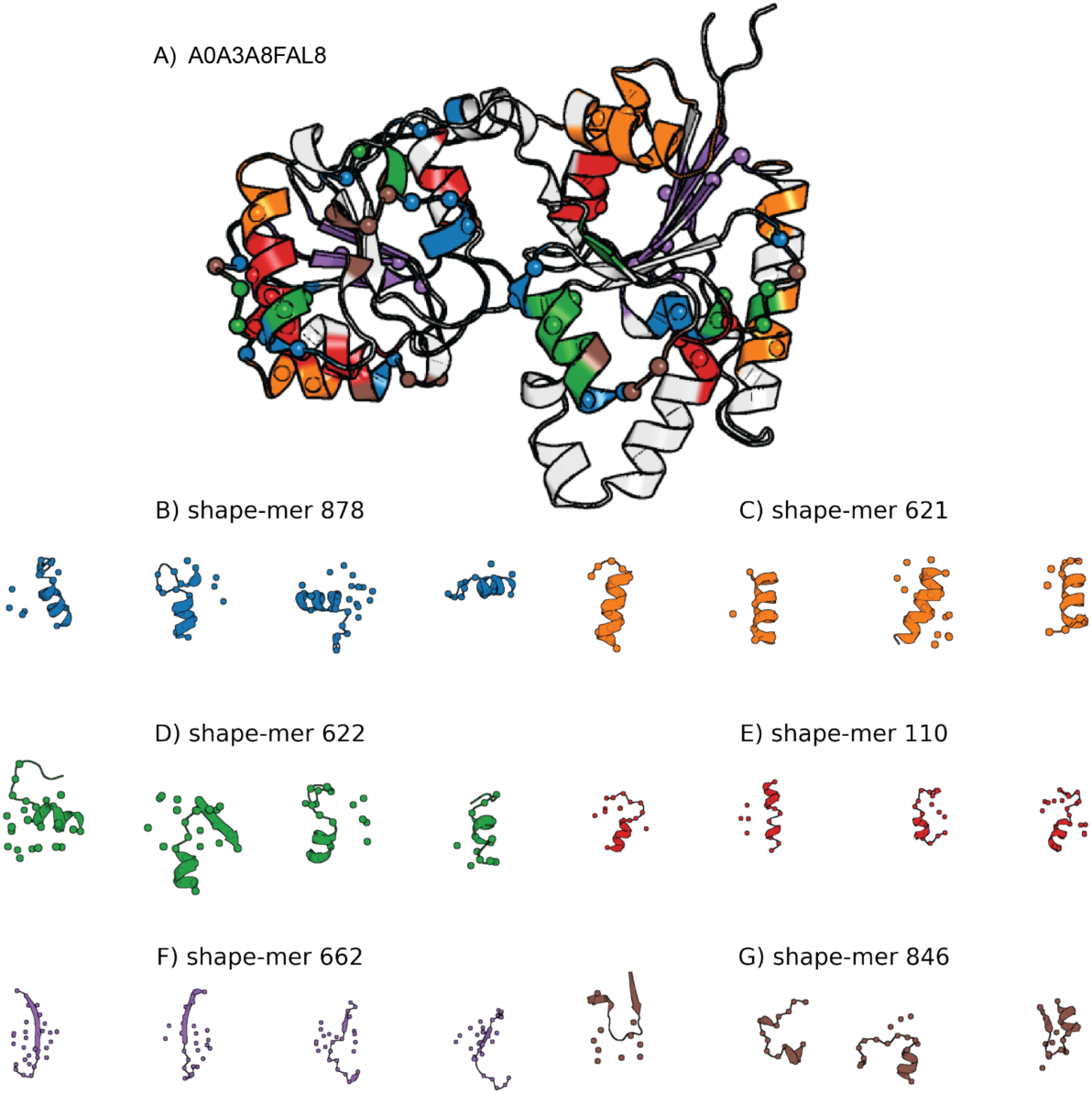
An example of substructure decomposition. (A) An example AlphaFold protein model with its 6 most common shape-mers highlighted in different colours. Spheres mark the shape-mer central residue and backbone atoms within 4Å are coloured. (B-G) Four random representatives of each selected shape-mer, obtained from CATH proteins with <20% sequence identity. Spheres depict positions within 8 residues in sequence and 10Å spatially from the central residue.

**Figure S5.**
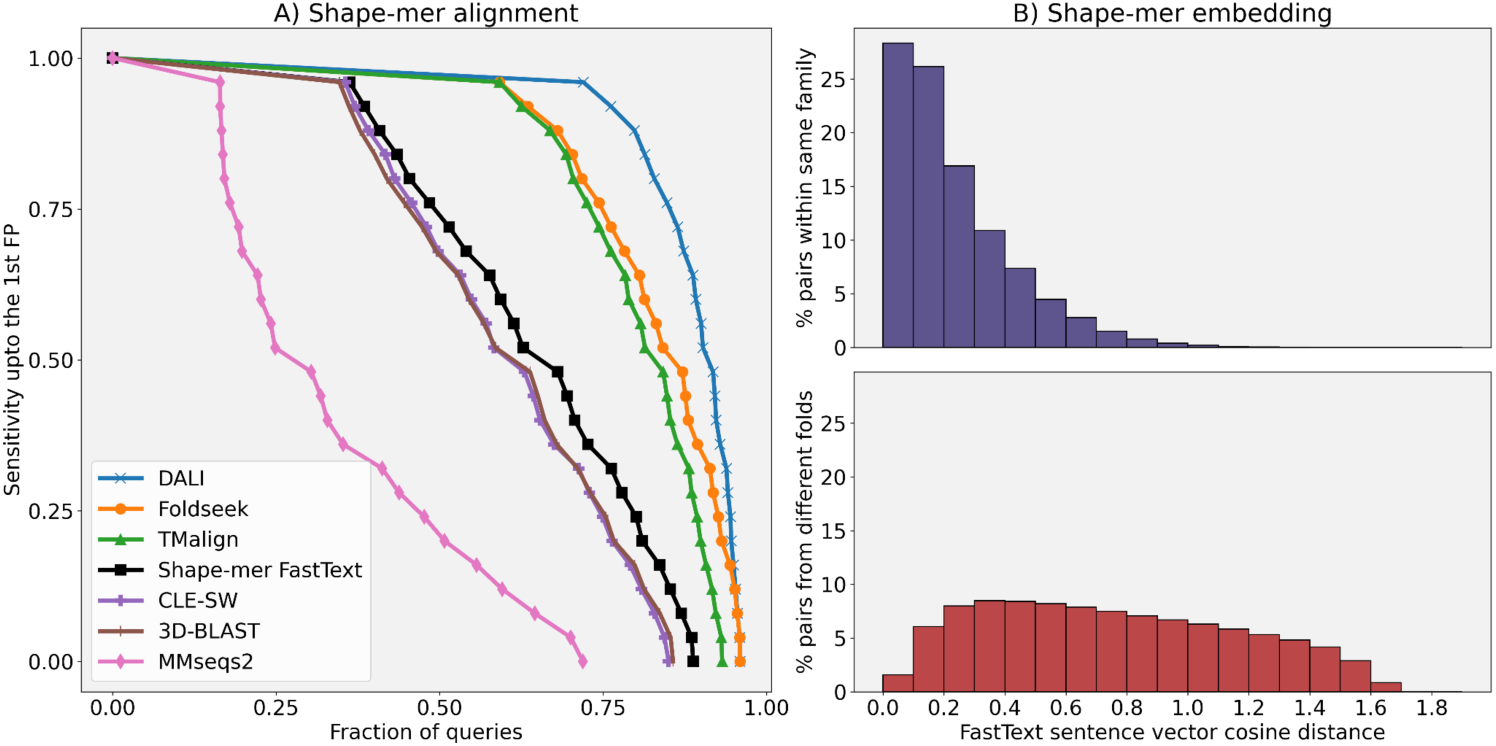
Shape-mer representations combined with FastText can discriminate between protein families. (A) Cumulative distributions of sensitivity for homology detection on the SCOPe40 database of single-domain structures. True positives (TPs) are matches within the same SCOPe family, false positives (FPs) are matches between different folds. Sensitivity is the area under the ROC curve up to the first FP. Results based on shape-mer FastText Smith-Waterman alignment are shown in black. Protein-level embedding distance measured as the cosine distance of FastText sentence vectors for proteins within the same SCOPe family (top) and from different SCOPe folds (bottom).

## SUPPLEMENTARY TABLES

**Supplementary Table S1.** List of 22 Pfams built from our data and incorporated into Pfam 36.0.

| Pfam ACC | Pfam ID | Description |
| --- | --- | --- |
| PF21619 | DUF6855 | Family of unknown function (DUF6855) |
| PF21651 | DUF6858 | Family of unknown function (DUF6858) |
| PF21738 | DJR_capsid | Double jelly roll capsid-like protein |
| PF21739 | DUF6866_N | Family of unknown function (DUF6866) N-terminal domain |
| PF21740 | DUF6866_C | Family of unknown function (DUF6866) C-terminal domain |
| PF21741 | DUF6867 | Domain of unknown function (DUF6867) |
| PF21742 | DUF6868 | Family of unknown function (DUF6868) |
| PF21746 | DUF6869 | Family of unknown function (DUF6869) |
| PF21747 | YpoC | YpoC-like protein |
| PF21757 | DUF6870 | Family of unknown function (DUF6870) |
| PF21758 | PAC_bac | Bacterial proteasome assembling chaperone-like protein |
| PF21777 | SDR-like | Short chain dehydrogenase-like protein |
| PF21778 | DUF6873 | Family of unknown function (DUF6873) |
| PF21779 | DUF6874 | Family of unknown function (DUF6874) |
| PF21780 | DUF6875 | Domain of unknown function (DUF6875) |
| PF21781 | DUF6876 | Family of unknown function (DUF6876) |
| PF21790 | OGG | Putative 8-oxoguanine DNA glycosylase OGG-like protein |
| PF21793 | DUF6877 | Family of unknown function (DUF6877) |
| PF21798 | DUF6878 | Family of unknown function (DUF6878) |
| PF21804 | Transposase_29 | Transposase protein |
| PF21805 | Imm5_like | Imm-5 like putative immunity protein |
| PF21806 | DUF6879 | Family of unknown function (DUF6879) |

**Supplementary File 1. Top 10 GO term predictions for the members of the clusters in the high-resolution sequence similarity network of component 159 (TumE) in figure 5A and their cognate antitoxin families, as predicted with DeepFRI**.

**Supplementary File 2. DNA fragments and DNA oligonucleotides used for plasmid construction during the experimental validation and characterisation of TumE and TumA**.

